## Supplementary material for "Large male proboscis monkeys have larger noses but smaller canines": SI 1 (from another version of this preprint: version 2, 2020-04-18)

### The nose is mightier than the tooth: larger male proboscis monkeys have smaller canines

**SI 1. List of subject information**

| Sex | Age/class | Nose size (cm, square root) | Body mass (kg, cube root) | Maxillary canine length (mm) | Note |
|---|---|---|---|---|---|
| Male 1 | Adult | 5.7 | 2.7 | 17.0 | |
| Male 2 | Adult | 5.7 | 2.8 | 20.5 | |
| Male 3 | Adult | 5.7 | 2.8 | 18.6 | |
| Male 4 | Young adult | 6.2 | 2.8 | 20.0 | |
| Male 5 | Old adult | 6.1 | 2.9 | 17.0 | |
| Male 6 | Adult | 5.9 | 2.7 | 22.1 | |
| Male 7 | Old adult | 5.8 | 2.8 | 17.2 | |
| Male 8 | Adult | 6.3 | 2.9 | 15.1 | |
| Male 9 | Adult | 6.0 | 2.7 | 20.0 | |
| Male 10 | Adult | 5.9 | 2.9 | 13.2 | |
| Male 11 | Adult | 5.3 | 2.8 | 14.2 | |
| Male 12 | Adult | 5.9 | 2.8 | 22.4 | |
| Male 13 | Adult | 6.0 | 2.8 | 20.3 | |
| Male 14 | Adult | 6.3 | 2.9 | 18.3 | |
| Male 15 | Adult | 6.2 | 2.8 | 12.2 | |
| Male 16 | Adult | 6.1 | 2.7 | 19.8 | |
| Male 17 | Adult | 6.1 | 2.7 | 24.2 | |
| Male 18 | Young adult | 5.6 | 2.7 | 20.0 | |
| Female 1 | Adult | 3.5 | 2.4 | 5.5 | |
| Female 2 | Adult | 3.2 | 2.3 | 7.6 | Pregnant |
| Female 3 | Adult | 3.5 | 2.4 | 8.2 | |
| Female 4 | Adult | 3.6 | 2.4 | 8.0 | Lactating |
| Female 5 | Adult | 3.5 | 2.4 | 9.3 | |
| Female 6 | Adult | 3.3 | 2.3 | 7.2 | |
| Female 7 | Young adult | 3.2 | 2.2 | 9.6 | |
| Female 8 | Adult | 3.1 | 2.2 | 8.3 | |
| Female 9 | Adult | 2.8 | 2.1 | 7.0 | Lactating |
| Female 10 | Adult | 3.7 | 2.2 | 8.6 | |
